# Analysis of SIV Spatiotemporal Dissemination Patterns in Rhesus Macaques During Early Rectal Transmission Demonstrates Systemic Infection Does Not Require Viral Local Amplification at the Entry Portal

**DOI:** 10.1101/2025.11.12.687929

**Authors:** Yilun Cheng, Jackson Chen, Miaoyun Zhao, Subhra Mandal, Saroj Chandra Lohani, Qin Liu, Mark Lewis, Ma Luo, Michael Gale, Qingsheng Li

## Abstract

Receptive anal sex is a dominant mode of HIV-1 (HIV) transmission, especially among men who have sex with men (MSM). However, the early events during HIV rectal transmission are not well understood and many questions remain, such as, does virus local amplification at the entry portal require for viral distal seeding? What is the spatiotemporal dissemination pattern across whole body? How do three viral forms, i.e., cell-free virus (V_CF_), cell-associated virus (V_CA_), and follicular dendritic cell-trapped virus (V_FDC_), evolve during early infections? To close this knowledge gap, we comprehensively examined rectum, draining lymph nodes (dLN) and distal lymph nodes (disLN), as well as non-lymphoid organs of brain, lungs, and liver of Indian rhesus macaques (RMs) at the early time points following intrarectal SIVmac251 inoculation (1, 2, 3, 4, 6, 10 14, 28 day post inoculation, dpi). Our findings demonstrated SIV rapidly disseminates to distant tissues and organs (<=1 dpi) and virus local amplification in rectum and dLN is not required for viral distal seeding for the establishment of systemic infection, viral forms shift from V_CF_ and V_CA_ to V_FDC_ accompanying from T cell zones into B cell follicles. Collectively, these findings indicate that an effective HIV vaccine needs to induce immune protection both locally at sites of viral entry and systemically.

## INTRODUCTION

Viral infection often begins with its crossing mucosal barrier and disseminating to distal sites through blood circulation, lymphatic system, neurological pathways along or in combination. A better understanding of virus dissemination patterns and the role of virus local amplification at the entry portal after exposure is critical for developing vaccines and pre- and post-exposure prophylaxis. Different viruses have distinct dissemination patterns after the exposure. For example, rabies virus dissemination follows a multi-stage process where viral local replication and amplification in muscle tissues after animal bite and scratch is critical for virus dissemination and widespread infection in the central nervous system (CNS) (1, 2) and Poliovirus local replication and amplification in the pharynx and the gastrointestinal tract has been demonstrated to be required for spreading to the CNS (3). However, the role of HIV local amplification at the portal of entry for systemic infection is unknown.

HIV infection remains a global health threat. About 39 million people are living with HIV worldwide. Despite the presence of a combined antiretroviral therapy and several preventive measures, approximately 1.3 million HIV new infections were still reported in 2022 (https://unaids.org/en). Receptive anal sex is a major mode of HIV transmission, especially in men who have sex with men (MSM) (4, 5), which accounted for 66% of all new HIV diagnoses in males in the United States (CDC, HIV surveillance report 2024, https://www.cdc.gov/hiv-data/nhss/hiv-diagnoses-deaths-and-prevalence-2025.html). Moreover, HIV vaccine development over the past four decades has made some progress, however, remains elusive. Thus, a better understanding of very early events of HIV mucosal transmission and viral spatiotemporal distribution patterns crossing viral entry site, draining and distal lymphoid organs and non-lymphoid organs of whole body, especially the role of viral local amplification at the portal of entry in the establishment of systemic infection, is needed for guiding HIV vaccine development (5–8). Nevertheless, for the convenience and ready availability of peripheral blood in clinical research, most investigations on HIV transmission have primarily depended on the continuous monitoring of peripheral blood samples (9–11). However, HIV target cells are mainly localized in mucosal and the secondary lymphoid tissues (LTs) and there is an eclipse phase between the point of initial infection to the time of viremia detection (12, 13). Due to ethical constraints in humans, in vivo data regarding viral entry and early dissemination patterns can only be studied using rhesus macaques (RMs) and simian immunodeficiency virus (SIV) model (14, 15). Nevertheless, previously published RM-SIV rectal transmission studies mainly focused on infection at the entry site and viral-host mucosal interactions (16–18). Ribeiro and colleagues found that SIV viral DNA was detected as early as 4 hours post inoculation (pi) in colic lymph nodes (LN), but not in axillary LN until 2 days post inoculation (dpi) (19), Sui and colleagues reported only 15% RMs had detectable viral RNA (vRNA) in the plasma as early as 4 dpi (20). Mair and colleagues reported that at 48 hours pi, infected cell foci consisting of only a few cells scattered throughout the anal and rectal tissues (21). We previously reported that SIV vRNA was only detectable in the rectum at 3 dpi but not at 1 and 2 dpi using radioactive S^35^ labeled SIV antisense probe in situ hybridization (ISH), which was much less sensitive than RNAscope ISH (RNAscope) (16). In short, there was a significant knowledge gap in the very early events of HIV rectal transmission, especially the role of virus local amplification for the systemic virus dissemination before the current study.

In this research, we comprehensively evaluated the viral spatiotemporal distribution patterns across whole body, the dynamic change of viral forms, i.e., cell-free virus (V_CF_), cell-associated virus (V_CA_), and follicular dendritic cell-trapped virus (V_FDC_), and the role of virus local amplification for the establishment of systemic infection using a highly sensitive RNAscope and a combination of CO-Detection by inDEXing (CODEX) immunostaining with RNAscope (Comb-CODEX-RNAscope). We found that virus local amplification at the portal of entry and draining lymph node (dLN) are not required for viral systemic infection; at the 1 dpi, vRNA were readily detected not only in the rectum and its dLN, but also in the distal LN (disLN) as well as non-lymphoid organs of liver, lungs and brain. These findings advance current concepts of early HIV rectal transmission by showing that rapid systemic dissemination occurs even in the absence of local viral amplification at the portal of entry, suggesting that an effective HIV vaccine needs to elicit robust immune protections at both the portal of viral entry and at the systemic level. An earlier version of this manuscript was posted to bioRxiv as a preprint (22).

## MATERIALS AND METHODS

### Animals and Ethics Statement

This report studied the samples from adult male RMs (*Macaca mulatta*) of Indian origin. All the animals were specific pathogen free (SPF, negative for HIV-2, SIV, type D retrovirus, and simian type D retrovirus 1) and without the protective major histocompatibility complex class I alleles Mamu A01, B01, and B17.

To delineate the earliest events and define the role of local viral amplification in the rectum and dLN for seeding distal sites and establishing systemic infection, RMs were intrarectally inoculated with SIVmac251 at the dose of 6,000 TCID_50_ (low-dose cohort) and were euthanized at 1, 2, 3, 6 dpi (n=4 per time point). The RM work was conducted at the Washington National Primate Research Center and was approved by the University of Washington Environmental Health and Safety Committee, the Occupational Health Administration, the Primate Center Research Review Committee, and the Institutional Animal Care and Use Committee (IACUC, protocol number 214207).

To study existing viral forms of V_CF_, V_CA_, and V_FDC_ in the LTs, RMs were intrarectally inoculated with SIVmac251 at a dose of 3.1 x 10^4^ TCID_50_ (high-dose cohort) and were euthanized at 3, 4, 6, 10, 14 and 28 dpi (n=3 for 3, 10, 14 and 28 dpi; n=4 for 4 and 6 dpi). Three RMs without virus inoculation were served as uninfected controls. The RM study was approved by the IACUC at the University of Nebraska-Lincoln (protocol number 559) and BIOQUAL, Inc. (protocol number 10-0000-01) and conducted at BIOQUAL, Inc. (Rockville, Maryland). All the tissues during necropsy were fixed with SafeFix II (Cat# 23-042600, Fisher Scientific), 4% paraformaldehyde or neutral-buffered formalin and embedded with paraffin as we previously reported (17, 23).

### RNAscope In Situ Hybridization (ISH)

RNAscope from the Advanced Cell Diagnostics (ACD) was performed according to our previously published method(24, 25). RNAscope® Probe-SIVmac239 anti-sense (SIV probe, Cat# 312811, ACD) was used and the signals were amplified and detected with RNAscope® 2.5 HD assay-Red kit (Cat# 322360, ACD). The RNAscope® negative control probe-DapB (Cat# 310043, ACD) was used as negative probe control and uninfected RM tissues were used as negative tissue control hybridized with SIV probe. Tissue sections on slides were digitized using our Aperio ScanScope into our image server and perform image quantification analysis using our Aperio Spectrum system as previously reported(25).

### Antibody-oligonucleotide conjugation

To find a RM reactive CD3 antibody for CODEX, an anti-human CD3 antibody in a carrier-free PBS solution (Clone#: SP162, Cat#: ab245731, Abcam) was conjugated with the barcode-oligonucleotide (Cat#: 5350002, Akoya) using CODEX conjugation kit (Cat#: 7000009, Akoya) following CODEX conjugation manual as we previously reported(26).

### Combination of CODEX immunostaining with RNAscope

(Comb-CODEX-RNAscope) CODEX is a cutting-edge multiplexed tissue imaging technique that enables simultaneous visualization of dozens of protein markers while preserving tissue architecture. Comb-CODEX-RNAscope enables detection of viral RNA via RNAscope and dozens of proteins. Comb-CODEX-RNAscope was performed according to our previously published method (26). CODEX and subsequent RNAscope approach and the combination of pretreatments of CODEX and RNAscope together at the beginning of Comb-CODEX-RNAscope were used(26). Briefly, tissue sections of 6-μm in thickness were mounted on poly-lysine-coated coverslips. The deparaffinization process began by heating samples in a 60°C incubator for 2 hours, followed by two 5-minute xylene washes. Samples were then rehydrated through a series of decreasing ethanol concentrations and diethyl pyrocarbonate (DEPC) water. Prior to initiating the CODEX cycling, tissues underwent pretreatment with 3% hydrogen peroxide and antigen retrieval in pH 6 citrate buffer (Sigma 21545).The retrieved tissues were then incubated with a DNA-barcoded antibody mixture for 3 hours at room temperature. Unbound antibodies were removed through three consecutive 2-minute PBS washes. Post-fixation occurred in two stages: first with 1.6% paraformaldehyde for 10 minutes at room temperature, followed by washing and a second fixation step using ice-cold methanol for 5 minutes. After additional washing, samples were stored in buffer at 4°C for up to five days or immediately processed in the CODEX instrument. Fluorescent reporters consisting of labeled oligonucleotide probes matching the DNA-barcoded antibodies were prepared in a 96-well plate according to the manufacturer specifications. The CODEX fluidics system and Keyence microscope operated under the control of the CODEX instrument manager and Keyence software following standard protocols. Upon completion of CODEX cycling, tissue-coverslips were transferred to the RNAscope workflow beginning at the protease plus treatment step. Following RNAscope completion, samples were counterstained with DAPI (1:2000 dilution) for 5 minutes at room temperature, washed, and returned to the CODEX instrument for RNAscope image acquisition. Integration of RNAscope data with CODEX images requires adding a blank cycle with Cy5-channel assignment for RNAscope detection. Final image data was processed using CODEX processor software after transferring to the appropriate file location.

### Data Analysis for the Comb-CODEX-RNAscope

After the processing of Comb-CODEX-RNAscope raw data with CODEX processor software, FlowJo v10 with CodexMAV extension was used to analyze data. Flow cytometry like gating strategy was used to determine cell populations. The statistical analysis and figures of cell populations and viral load dissemination change were made with Graphpad Prism 9.5.

### Spatial Cluster Analysis of vRNA Distribution

Spatial analysis of vRNA distribution was performed using QuPath software (https://qupath.github.io/). Prior to analysis, the metadata for all whole tissue section images was manually verified to ensure correct pixel size calibration. A pixel classifier was first trained to segment tissue regions by distinguishing tissue from the white background. To separate the stains, color deconvolution was performed. Initial stain vectors for nuclear stains with Hematoxylin and vRNA signal of Fast Red were identified using the ‘Estimate Stain Vectors’ command, followed by manual adjustment to ensure optimal stain separation. Following segmentation, vRNA-positive signals were identified using the ‘Positive Cell Detection’ command. The analysis was run at a requested pixel size of 0.5 µm/pixel. Key parameters were set as follows: the background radius was 5 µm, the minimum and maximum detection areas were set to 0.5 µm² and 5.0 µm², respectively, and cell expansion was set to 0 µm. To identify spatial clusters of vRNA-positive cells, the Density-Based Spatial Clustering of Applications with Noise (DBSCAN) algorithm was subsequently applied(27). The algorithm’s parameters were set with an epsilon (maximum search radius) of 50 µm and a minimum of 3 points (minPts) required to form a dense cluster.

## RESULTS

### SIV fast disseminates to dLN and disLN as well as non-lymphoid organs of lungs, liver and brain during very early rectal transmission

To assess SIV spatiotemporal dissemination after rectal transmission, RMs were intrarectally inoculated with SIVmac251 at the dose of 6,000 TCID_50_ (low-dose cohort) and euthanized at 1, 2, 3, 6 dpi (n=4 per time point). Additionally, three RMs without SIV inoculation were euthanized to serve as uninfected controls. A comprehensive necropsy was performed on each animal to collect and fix samples, including rectum, dLN (mesenteric), disLN (axillary and mandibular), spleen, and non-lymphoid organs of liver, lungs, and brain (Fig.1). Peripheral blood was collected before and after virus inoculation and plasma was stored at −80C for SIV vRNA quantification using qRT-PCR.

**Figure 1.**
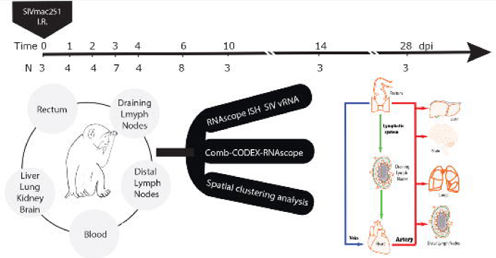
Experimental design and sampling timeline of rhesus macaques following rectal SIVmac251 inoculation. Rhesus macaques were sacrificed at 1–28 days post inoculation (dpi), as indicated on the timeline with animal numbers per time point, for collection of rectum, draining lymph nodes (dLN), distal lymph nodes (DisLN), blood, and distal organs. Tissues were analyzed by RNAscope for SIV viral RNA (vRNA), combined CODEXIZRNAscope multiplex imaging, and spatial clustering, and the schematic illustrates proposed viral spread from rectal mucosa through the lymphatic and circulation system to systemic organs.

To detect SIV vRNA in tissues, RNAscope with SIV probe was performed, of which the detection sensitivity is a single copy of vRNA. To ensure the specificity of vRNA detection by RNAscope, we included the tissues of SIV-infected RMs that were hybridized with the bacterial DapB probe and the tissues of SIV-uninfected RMs that were hybridized with SIV probe as negative control. These controls collectively confirm the specificity of the detected vRNA signals. Background signals were exceedingly rare and, when present, appeared weak, diffuse, and predominantly localized to tissue edges, consistent with edge effects. In contrast, vRNA signal was defined by its strong intensity, discrete punctate appearance, and localization within morphologically intact cells or anatomically relevant tissue regions (Fig. S1). Moreover, we validated vRNA detection specificity using RNAscope by incorporating RNase digestion in adjacent sections form SIV-infected RMs, which abolished vRNA signals and thereby confirmed that the detected signals were RNA-specific. Thus, the observed vRNA signals represent true vRNA rather than nonspecific background. On 1 dpi, vRNA was readily detected in the rectal and dLN of all RMs, and importantly, also in the disLN (axillary and mandibular) (Figs. 2 & S2) and non-lymphoid organs of lungs, liver and frontal cortex and basal ganglion of brain (Fig. 3, Table S1), demonstrating that at 1 dpi, SIV already disseminated from the portal of rectal entry to dLN, disLN, distal non-lymphoid organs. Of note, the frequency and abundance of detected vRNA signals were low, manifesting as a single dot, which morphologically resembles a single virion derived signal. At 2, 3 (Figs 4 & S3 & Table S1) and 6 dpi (Fig. 5A & B), vRNA signals were readily detected in all the tissues that we examined, including rectal, dLN, disLN, and distal non-lymphoid organs in all RMs. Noteworthy, as we previously reported plasma viral load (pVL) at 1, 2 and 3 dpi was undetectable in all RMs and only became detectable at 6 dpi using a highly sensitive qRT-PCR (limit of detection, 1 copy/ml) (28). To our knowledge, this is the first study that comprehensively investigated spatiotemporal viral dissemination across tissues and organs in such an early timeframe.

**Figure 2.**
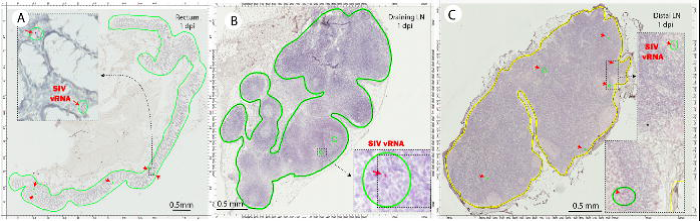
Detection of SIV vRNA in rectum, dLN and disLN at 1 dpi. Representative micrographs show SIV vRNA in rectal mucosa (**A**), mesenteric dLN (**B**), and mandibular disLN (**C**) at 1 dpi from rhesus macaques infected with SIVmac239. Tissue regions of interest are outlined (rectum and dLN in green; disLN in yellow), and individual vRNAIZpositive cells appear as distinct red punctate signals detected using SIVmac239 antisense probes and Fast Red chromogen, with hematoxylin counterstaining of cell nuclei (blue). Red arrows and green circles highlight discrete vRNA signals within each tissue section. HigherIZmagnification inset images from the boxed areas in each panel illustrate representative SIV vRNA–positive cells and their spatial distribution within the rectal laminar propria and lymph node parenchyma. Scale bar, 50 mm.

**Figure 3.**
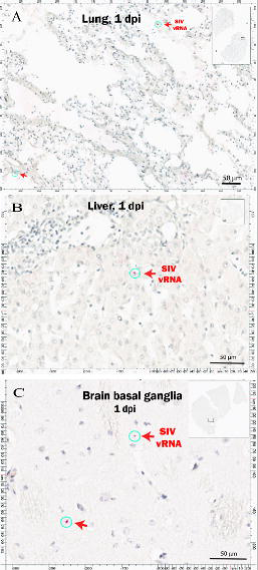
Detection of SIV vRNA in non-lymphoid organs of lungs, liver and brain at 1 dpi. Representative micrographs shows SIV vRNA in lung (**A**), liver (**B**), and brain basal ganglia (**C**) at 1 dpi from rhesus macaques infected with SIVmac239. SIV vRNA appear as distinct red punctate signals detected using RNAscope with SIVmac239 antisense probes and Fast Red chromogen, with hematoxylin counterstaining of cell nuclei (blue). Insets depict low-magnification images with boxed regions corresponding to the higher-magnification fields shown. Red arrows and green circles highlight SIV vRNA–positive cells. Scale bar, 50 μm.

**Figure 4.**
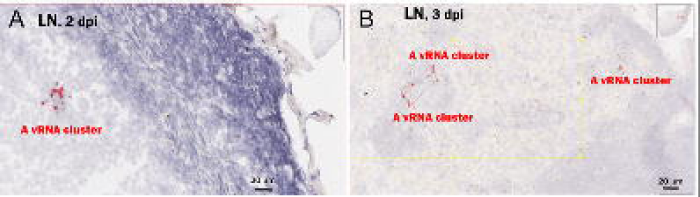
Spatial clustering of SIV vRNA in lymph node tissues on 2 and 3 dpi. Representative micrographs show SIV vRNA clusters within dLN tissues collected from rhesus macaques at 2 and 3 dpi. vRNA clusters were identified through implementation of the Density-Based Spatial Clustering of Applications with Noise (DBSCAN) algorithm within QuPath image analysis software. A vRNA cluster is defined as ≥3 individual vRNA signals within 50 μm diameter regions. vRNA was detected using RNAscope. Higher magnification insets from the boxed regions of each tissue section are displayed.

**Figure 5.**
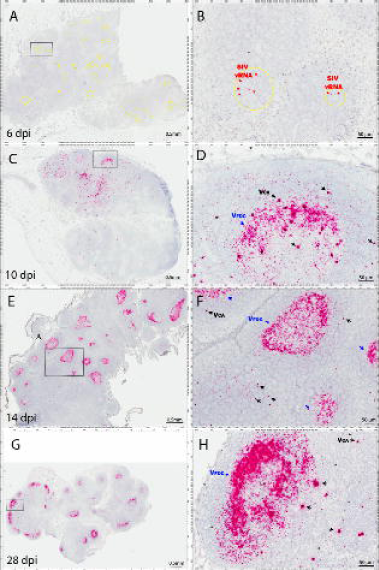
Temporal and spatial dynamics of three distinct viral forms in lymph node tissues during acute SIV rectal infection. Representative micrographs show three distinct viral forms, i.e., cell-free viruses (V_CF_), cell-associated viruses (V_CA_), and follicular dendritic cell-trapped viruses (V_FDC_) in lymph node tissues during early rectal infection. At 6 dpi ( **A, B**), vRNA was exclusively detected as V_CA_ and V_CF_ populations, with V_CF_ appearing as discrete signals external to cellular boundaries. By 10 dpi (**C, D**), V_FDC_ became detectable for the first time, characterized by diffuse vRNA signal patterns in the B-cell follicles. The V_FDC_ population emerged as the predominant viral form at 14 dpi (**E, F**) and 28 dpi (**G, H)**. Higher magnification images on the right side of each panel provide detailed visualization of the boxed regions from the corresponding left-side overview images. Scale bars, 0.5 mm (left) and 50 μm (right).

### Virus Local Amplification at the Portal Entry Is Not Required for Establishing Systemic Dissemination

To determine whether viral distal seeding is the consequence of viral local amplification at the portal of entry and dLN, we ran a spatial cluster program to quantify virus amplification clustering. A viral cluster is defined as there are 3 or more distinct virions derived from vRNA signals in a closely adjacent area of smaller than 50 μm in radius. As the diameter of CD4 T cells is generally 5–10 µm (29) and three or more of virion in such as small area indicate viral local amplification. On 1 dpi, when vRNA was detected in disLN, spleen and non-lymphoid organs of liver, lungs and brain, we did not find any viral clusters in these tissues in all RMs (Figs. 2 & S2), indicating virus local amplification at portal rectal entry and dLN is not essential for virus distal seeding. In contrast, we detected viral clusters in both dLN and disLN at 2, 3, and 6 dpi (Figs. 3 & 4A-B) when SIV was already widely disseminated to distal tissues and organs, indicating vRNA replication and amplification across the body after establishing the systemic dissemination. To compare the proportions of RMs with viral cluster between 1 dpi versus 2, 3, and 6 dpi, Fisher’s exact test was used and the test result indicates that there is a statistically significant difference on proportions of animals with viral cluster between 1 versus 2, 3, 6 dpi respectively (0/4 on 1 dpi versus 4/4 on 2, 3, 6 dpi, p-value=0.0286, two-sided p-value). To our knowledge, this is the first study showed evidence that SIV local amplification at the portal of entry is not required for the establishment of systemic infection.

### The Spatiotemporal Distribution Patterns of Three Different Viral Forms and Virally Infected CD4 and Macrophage during Early Rectal Transmission

Three viral forms, i.e., V_CF_, V_CA_, and V_FDC_, can coexist in vivo in LTs during HIV infection. Different viral forms play distinct roles in the establishment and maintenance of infection and pathogenesis. To examine viral forms during early infection, RMs were inoculated with SIVmac251 at a dose of 3.1× 10^4^ TCID_50_ (high-dose cohort) and euthanized at 3, 4, 6, 10, 14, and 28 dpi (n = 3-4 per time point). This cohort overlapped with the low-dose cohort at 3 and 6 dpi. To assess potential dose-dependent effects, we performed cohort-stratified comparisons at these two shared time points. No significant differences were observed between cohorts in the distribution of vRNA or spatial dissemination patterns. Viral clusters were readily detectable at 3 and 6 dpi in both cohorts, indicating that inoculum dose did not measurably alter early dissemination dynamics in our study. These findings are consistent with prior SIV challenge studies demonstrating that, although inoculum dose strongly influences acquisition probability, established infections across a typical dose range rapidly converge in systemic dissemination kinetics(30, 31).

To quantitatively examine the spatial distribution patterns of different viral forms and virally infected cells in LTs during early rectal infection, we used the Comb-CODEX-RNAscope, a multiplexed proteins and SIV vRNA co-detection method(26) to visualize and quantify vRNA and multiple immune cell markers simultaneously. A cocktail of antibodies to CD3, CD4, CD68, CD20, CD21, CD31, Ki67, HLA-Dr, and IDO were used to identify immune cell types and immune activation state in combination with vRNA detection. Fig. 6 shows representative images for co-detection of vRNA and various immune markers. We first quantified total vRNA disregarding virus forms in T-cell zones versus B-cell follicles in LTs during early infection by using the B-cell follicle marker CD20 to gate CD20⁺ B-cell follicles and CD20⁻ T-cell zones. vRNA in vivo growth curve (1, 2, 3, 6, 10, 14, 28 dpi, n=3-4/time point) in T-cell zones peaked at 10 dpi and then declined, while vRNA in B-cell follicles significantly increased at 10 dpi and maintained at high level till 28 dpi (Fig. 7A-C). We also quantified CD4 T cells and immune activation markers in the LN tissues of uninfected and infected RMS at 6, 10, and 28 dpi. There was a significant CD4 T cell decline and a significantly increased expression of immune activation markers of Ki67, HLA-Dr (Fig. S4) as well as immune suppression protein marker of Indoleamine 2,3-dioxygenase 1 (IDO1) that may counteract the overall immune activation induced by SIV infection (Fig. S4).

**Figure 6.**
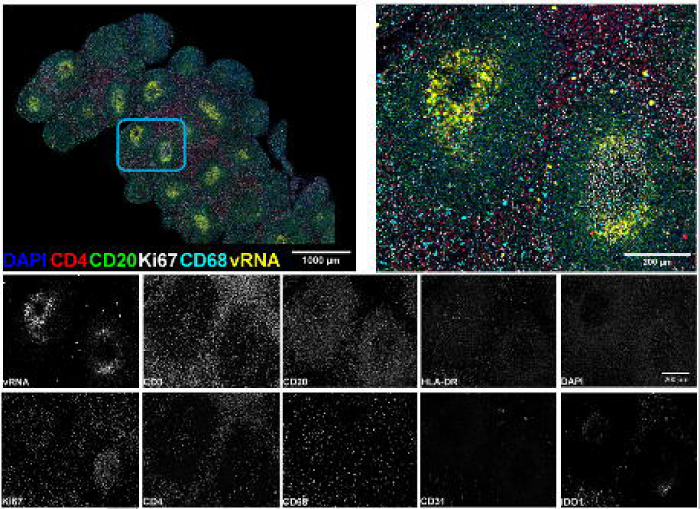
Representative Comb-CODEX–RNAscope imaging of lymph node tissues from an SIV-infected rhesus macaque. The upper left panel shows a stitched overview of an entire lymph node cross-section (28 dpi, animal Rh5429) with multiplex fluorescence for DAPI (nuclei, blue), CD4 T cells (red), CD20 B cells (green), Ki67 proliferating cells (white), CD68 myeloid cells (cyan), and vRNA yellow), illustrating the spatial organization of B-cell follicles and paracortical regions (scale bar, 1000 µm). The cyan box denotes the area enlarged in the upper right panel, which provides a higher-magnification view of Comb-CODEX–RNAscope signals highlighting vRNA-positive foci within and adjacent to B-cell follicles and the surrounding T-cell zone (scale bar, 200 µm). The lower panels display individual grayscale channels from the same high-magnification field for vRNA, CD3, CD20, HLA-DR, DAPI, Ki67, CD4, CD68, CD31, and IDO1, enabling visualization of single-marker distributions and their relationship to sites of viral RNA signal (scale bar, 200 µm).

**Fig 7.**
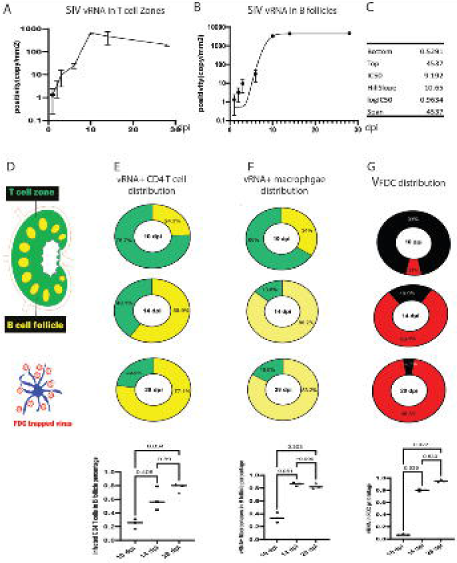
The spatiotemporal distribution patterns of virally infected cells and viral forms in lymph node tissues during early rectal transmission. SIV vRNA and immune cell markers were detected using Comb-CODEX-RNAscope and the spatial distribution patterns of vRNA+ cells in T-cell zones versus B-cell follicles of mesenteric LN tissues over time were quantified by using CODEX processor software. In vivo viral growth curves show the density of SIV vRNA–positive signals (copies/mm²) at 1, 2, 3, 6, 10, 14, and 28 dpi in T-cell zones (**A**) and B-cell follicles (**B**), with nonlinear fitting parameters shown for B-cell follicular viral load (**C**). Schematic representation of lymph node architecture, illustrating T-cell zones, B-cell follicles, and V_FDC_ (**D**). (**E-G**) Donut plots depict the percentage of SIV vRNA+ CD4 T cells (**E**) and vRNA+ macrophages (**F**) localized in T-cell zones (green) versus B-cell follicles (yellow) at 10, 14, and 28 dpi. (**G**) Donut plots show the proportion of V_FDC_ (red) within total vRNA signals (black plus red) in B-cell follicles at 10, 14, and 28 dpi. Bottom panels: quantification of the fraction of vRNA+ CD4 T cells (left), vRNA+ macrophages (middle), and V_FDC_ signals (right) present in B-cell follicles at 10, 14, and 28 dpi; horizontal bars indicate group medians, and P values are from non-parametric pairwise comparisons (10, 14, and 28 dpi).

In the very early infection (≤6 dpi), all the vRNA we detected are either V_CA_ or V_CF_ (Fig 5A & B). V_FDC_ was first detected at 10 dpi and became dominant viral form in LTs at 14 and 28 dpi (Fig. 5C-H). As LTs at 10,14 and 28 dpi contained all three viral forms corresponding to peak viremia (10-14 dpi) and the subsequent viral set point (28 dpi), which are critical for understanding HIV pathogenesis and the dynamic equilibrium of virus–host interactions, we thus focused on 10, 14, and 28 dpi samples to study the distribution patterns of three viral forms and virally infected CD4 and macrophages in LTs. Using the Comb-CODEX-RNAscope, we found that CD4 T cells are the primary cell type supporting productive viral replication (vRNA+), making up greater than 90% of infected cells, with macrophages constituting a minor fraction below 10% (data not show). We further quantified v*RNA*+ CD4 T cells and macrophages in T cell zones and B cell follicles of LN tissues (Fig. 7E). We first identified virally infected CD4 (CD4+ & vRNA+) or macrophage (CD68+ & vRNA+) by combining two channels of signals, then used B-cell follicle marker CD20 as a gate to differentiate virally infected cells in B-cell follicles versus T-cell zones. The population of virally infected CD4 T cells distribution patterns shifted from mainly in T cell zones at 10 dpi (75.7%) to mainly in B-cell follicles at 14 (T cell zones: 40.1%) and 28 dpi (T cell zones: 22.9%, Fig. 7E). We used two-sided Mann–Whitney U tests for pairwise comparisons and the Kruskal–Wallis test with post hoc correction for multiple comparisons to evaluate the distribution of virally infected CD4 T cells between T cell zones and B cell follicles across time points (10, 14, and 28 dpi). There is a significant increase in the accumulation of virally infected CD4 T cells within B cell follicles at 28 dpi compared with 10 dpi (P = 0.034; Fig. 7E). The distribution of virally infected macrophages within B cell follicles relative to T cell zones increased from 10 dpi, 14 dpi to 28 dpi (T cell zones: 66%, 13.6%, and 16.8%, respectively; Fig. 7F); however, these differences were not statistically significant (Fig. 7F). The V_FDC_ viral form constituted a small fraction of the total viral load at 10 dpi (17%) but increased markedly to 80.1% at 14 dpi and 95.3% at 28 dpi (Fig. 7G). This represents a significant increase in V_FDC_ at 28 dpi compared to 10 dpi (P = 0.022; Fig. 7G).

## Discussion

The period of time between HIV initial exposure at the mucosal site and the virus spreads throughout the body represents a critical window of time during which interventions could be most effective (32, 33). Historically using less sensitive viral detection methods, it was believed that a prolonged local viral amplification phase for small founder populations of infected cells near the portal of entry is essential for the establishment of systemic infection (7, 34–39). Deleage and colleagues, using barcoded-virus experiments and high-resolution mapping, indicate that multiple microfoci within the female genital tract appear very early (hours–days) preceded lymphatic dissemination, also before expansion and spread to distal sites (37). Historically, tissue-level studies using radiolabeled ISH and immunohistology showed high LN viral loads early after a few days of infection (24), and more modern RNAscope/IF-FISH approaches have increased sensitivity and allowed cell-type colocalization (26, 40–42). A convergence of data from rigorous necropsy studies in RM rectal inoculation models now reveal a dramatically accelerated timeline of systemic infection (43, 44). Results from Ribeiro and colleagues showed viral DNA or infectious virus in mesenteric dLN and local tissues as early as 4 hours after rectal exposure, consistent with very rapid trans-epithelial transport or virus-bearing cell trafficking. Within 2-3 days, robust replication was evident in systemic lymph nodes and the gut-associated lymphoid tissue (GALT), which rapidly becomes a major site of viral amplification (44). When studying the mechanisms of rapid viral spread during acute infection stage, some papers demonstrate that cell-associated inoculation can seed tissues faster because infected cells migrating from the mucosa into lymphatic channels appear to be one vehicle for early dissemination (37, 45). However, the whole-body analysis of viral spatiotemporal distribution patterns following early rectal transmission is unknown until this study.

Using the low-dose cohort of RMs that were intrarectally inoculated with SIVmac251 at the dose of 6,000 TCID_50_ and were euthanized at 1, 2, 3, 6 dpi, we comprehensively analyzed vRNA spatial distribution in rectal, dLN and disLN and nonlymphoid organs of lungs, liver and brain. We unambiguously demonstrated that SIV rapidly disseminated to rectum, dLN, disLN, spleen, liver, lungs and brain at 1 dpi and thereafter in all RMs using the ultra-sensitive RNAscope that enables us to detect single SIV virion (Fig. 2, 3, 4, 5, S2-3). We recognize that contextualizing the viral dose used in our study with respect to the number of infectious virions present in human semen is important, although such comparisons remain inherently complex. Estimates of infectious HIV particles in human ejaculate vary widely depending on viral load, stage of infection, and assay methodology. While total viral RNA copies in semen can reach 10^5^–10^7^ per mL(46, 47). Moreover, transmission across the rectal mucosa is inefficient, with only a small fraction of virions successfully initiating infection. Thus, although our inoculum may exceed the number of transmitted founder virions, it falls within the range commonly used in NHP models to overcome stochastic barriers to mucosal infection. Meanwhile, it remains inherently challenging to fully distinguish between the virus distal spread driven by infected cells or cell-free virus through blood circulation versus lymphatic system or in combination. With that said, the earliest detected vRNA in this study is consistent with early systemic dissemination events, where both blood and lymphatic circulation are likely involved (Fig. 1). First, the vRNA-positive puncta are not uniformly distributed but instead appear in discrete anatomical locations within tissues, rather than showing a diffuse vascular-associated pattern that might be expected from passive blood-borne virions. Second, the detection of vRNA in disLN and non-lymphoid organs of liver, lungs and brain at 1 dpi using RNAscope occurred in the absence of measurable systemic viremia, arguing against passive blood-borne carriage as the sole explanation and supporting early tissue seeding. Third, the frequency of these events is low and comparable across animals on the same dpi, arguing against widespread nonspecific deposition of circulating viruses. Fourth, in LTs, these signals are observed in proximity to target CD4 T cell-rich microenvironments, which are consistent with previously described early viral encounter sites. Moreover, our findings align with the recent research conducted by Whitney and colleagues (14) that examined the effects of initiating ART at 6 hours and 1 day following intrarectal SIVmac251 infection. Following cessation of the 6-month ART regimen, viral rebound rates were 0% in animals that began treatment at 6 hours post-infection and 20% in those that started treatment at 1 dpi. Regarding virus CNS dissemination, prior work in HIV and SIV models supports a predominant “Trojan horse” mechanism in which infected monocytes and CD4 T cells cross the blood–brain barrier and differentiate into perivascular macrophages, thereby introducing virus into the CNS(48, 49), however, our present study was not designed to dissect the relative contribution of specific neuroanatomical routes to brain dissemination.

To determine the role of viral local amplification at the portal of rectal entry and dLN for a distal seeding for the establishment of systemic infection, we performed a comprehensive analysis of vRNA spatial distribution in tissues from RMs in the low-dose cohort. We hypothesized that if a local viral amplification plays a crucial role in establishing distant viral seeding during initial systemic dissemination (1 dpi), then the evidence of viral local replication and expansion should be detectable within rectal and dLN tissues. We tested this hypothesis by using ultra-sensitive RNAscope and extensively examining vRNA in multiple sections of rectum, dLN, disLN and non-lymphoid organs, however, we did not detect any viral cluster in rectum and dLN, indicating that viral local amplification is not essential for virus distal seeding and establishment systemic infection. Our choice of ≥3 vRNA signals within a 50 µm radius was intended as a conservative operational definition to distinguish likely local amplification from stochastic proximity of independently arriving virions. The radius of 50 µm approximates the spatial scale of immediate cellular neighborhoods in lymphoid tissue (roughly 5-8 lymphocyte distance), where short-range cell-to-cell spread is most likely to occur. Requiring ≥3 signals reduces the probability that a “cluster” reflects the chance juxtaposition of two unrelated virions. Moreover, at 2, 3 and 6 dpi, viral clusters consistently contained ≥3 discrete vRNA signals within the same radius and there are multiple clusters in a single tissue section. These results support the interpretation that the absence of detectable clusters at 1 dpi is not solely a consequence of thresholding but is consistent with a lack of detectable local amplification at this stage.

Understanding the spatial distribution pattern of virally infected cells in T cell zones versus B cell follicles of LTs is essential for elucidating early virus–host interactions and their immunopathogenic consequences and informing strategies to prevent systemic dissemination. Using the Comb-CODEX-RNAscope platform in this study, we simultaneously visualized vRNA alongside immune cell markers, enabling high-resolution mapping of infected cell types, viral forms and their distribution within T cell zones versus B cell follicles in LTs. We found that the distribution of virally infected CD4 T cells shifted from predominantly within T cell zones at 10 dpi to primarily within B cell follicles at 14 and 28 dpi (Fig. 7E). A similar redistribution was observed for virally infected macrophages (Fig. 7F), paralleling the transition of vRNA localization from T cell zones at 10 dpi to B cell follicles at 14 and 28 dpi, particularly in V_FDC_ viral form (Fig. 7G).

FDCs reside in the light zone of germinal center in the B cell follicles and express complement receptor 2 (CR2/CD21), which capture and retain complement-opsonized HIV virions forming immune complexes that persist within germinal centers (50). Using Comb-CODEX-RNAscope, we simultaneous visualization of SIV vRNA, FDCs, and B cell follicles in SIV infected RM LN. Figure S5 shows the colocalization of VFDC with the CD21+ FDC network within CD20+ B cell follicles (Fig. S5). V_FDC_ has been demonstrated to be infectious to migrating CD4 T follicular help cells (Tfh) and play an important role in the HIV pathogenesis(51). The B-cell follicles in LTs are unique anatomically and functionally, as there is a low frequency of CD8 T cells and a high abundance of infectious V_FDC_, thus B follicles may be more favorable niche for viruses to survival by avoiding host’s immune attack and uninfected CD4 T cell and macrophages get infected by V_FDC_. Thus, the redistribution of virally infected cells into B cell follicles, along with increased V_FDC_ at 28 dpi, represents a mechanism of viral escape from host immune defenses.

We would like to point out that the sample size at each time point is relatively limited, which may reduce the statistical power to detect subtle differences. This constraint is inherent to nonhuman primate studies due to ethical and logistical considerations. Therefore, the findings should be interpreted with caution and considered preliminary rather than definitive.

In summary, using the RM-SIV rectal transmission model, combined with highly sensitive RNAscope and integrated Comb-CODEX–RNAscope analyses, and comprehensive examination of the rectum, dLN and disLN, as well as non-lymphoid organs including liver, lungs, and brain, we demonstrate that SIV rectal transmission does not follow a staged dissemination model. Instead, local viral amplification at the portal of entry is not required for the establishment of systemic infection. These findings indicate that an effective HIV vaccine must confer robust protection not only at mucosal entry sites but also at the systemic level.

---

**Figure S1. Validation of RNAscope specificity for in situ detection of SIV vRNA in rhesus macaque tissues.** (**A**) Representative micrograph showing lymph node section from an SIV-uninfected rhesus macaque that was hybridized with the SIV antisense probe shows absence of specific vRNA signal. (**B**) Lymph node section from an SIV-infected rhesus macaque at 14 dpi that was hybridized with the bacterial *dapB* probe as a negative control, demonstrates minimal background staining. (**C**) Lymph node section from an SIV-infected rhesus macaque at 14 dpi that was hybridized with the SIV antisense probe, shows abundant, discrete punctate vRNA signals localized within morphologically intact cells and anatomically relevant tissue regions. Scale bar, 0.5 mm.

**Figure S2. Detection of SIV vRNA in rectum, dLN, and disLN at 1 dpi.** Representative micrographs from RMs infected with SIVmac239, showing SIV vRNA-positive cells in rectal mucosa (left panels, A1–A4), mesenteric dLN (middle panels), and mandibular disLN (right panels) at 1 dpi. Tissue regions of interest are outlined (rectum and draining lymph nodes in green; distal lymph nodes in yellow), and individual vRNA-positive cells appear as discrete red punctate signals detected using SIVmac239 antisense probes and Fast Red chromogen, with hematoxylin counterstaining of cell nuclei (blue). Red arrows and green circles highlight discrete vRNA signals within each tissue section. Higher-magnification inset images from boxed areas in each panel illustrate representative SIV vRNA–positive cells and their spatial distribution within the rectal lamina propria and lymph node parenchyma. Scale bar, 0.5 mm.

**Figure S3. Detection of SIV vRNA in rectum, dLN, and disLN at 2and 3 dpi.** Representative micrographs show SIV vRNA in mesenteric dLN and mandibular disLN from rhesus macaques at 2 dpi (animal B1 & B4) and 3 dpi (animal C1 &C2). Green outlines indicate tissue borders and green circles denote discrete vRNA-positive foci detected with RNAscope using SIVmac239 antisense probes and Fast Red chromogen, with hematoxylin counterstaining of cell nuclei. Higher-magnification views of the boxed regions (red rectangles) are displayed adjacent to each corresponding low-magnification image to illustrate representative clusters of vRNA-positive cells. Scale bars in the higher-magnification panels represent 100 μm.

**Figure S4. Immune activation and CD4 T cell decline in the lymph node tissues during early SIV rectal infection.** Immune cell markers were detected using Comb-CODEX-RNAscope and quantified in the lymph node tissues of rhesus macaques infected with SIV at 6 10 and 28 dpi as well as uninfected controls (n=3/time point). SIV infection was associated with a significant decline in CD4 T cells, measured as the CD4/CD3 ratio, alongside increased expression of immune activation markers Ki67 and HLA-DR, and the immunosuppressive marker indoleamine 2,3-dioxygenase 1 (IDO1).

**Figure S5. Spatial distribution of V_FDC_ relative to CD21 and CD20 within B-cell follicles.** Representative micrographs show vRNA signal in the viral form of V_FDC_ (**A**), CD21 (**B**), and CD20 (**C**) in lymph node tissues of SIV infected rhesus macaques at 28 dpi (animal # Rh5429) detected using Comb-CODEX-RNAscope. A merged multiplex fluorescence image is presented in (**D**), showing vRNA (white), CD20⁺ B cells (green), and CD21⁺ FDC networks (blue

