## Supplementary figures and images for "Analysis of SIV Spatiotemporal Dissemination Patterns in Rhesus Macaques During Early Rectal Transmission Demonstrates Systemic Infection Does Not Require Viral Local Amplification at the Entry Portal"

### Supplemental Figure 1

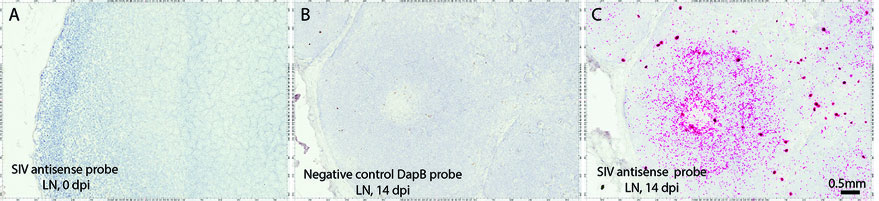

### Supplemental Figure 2

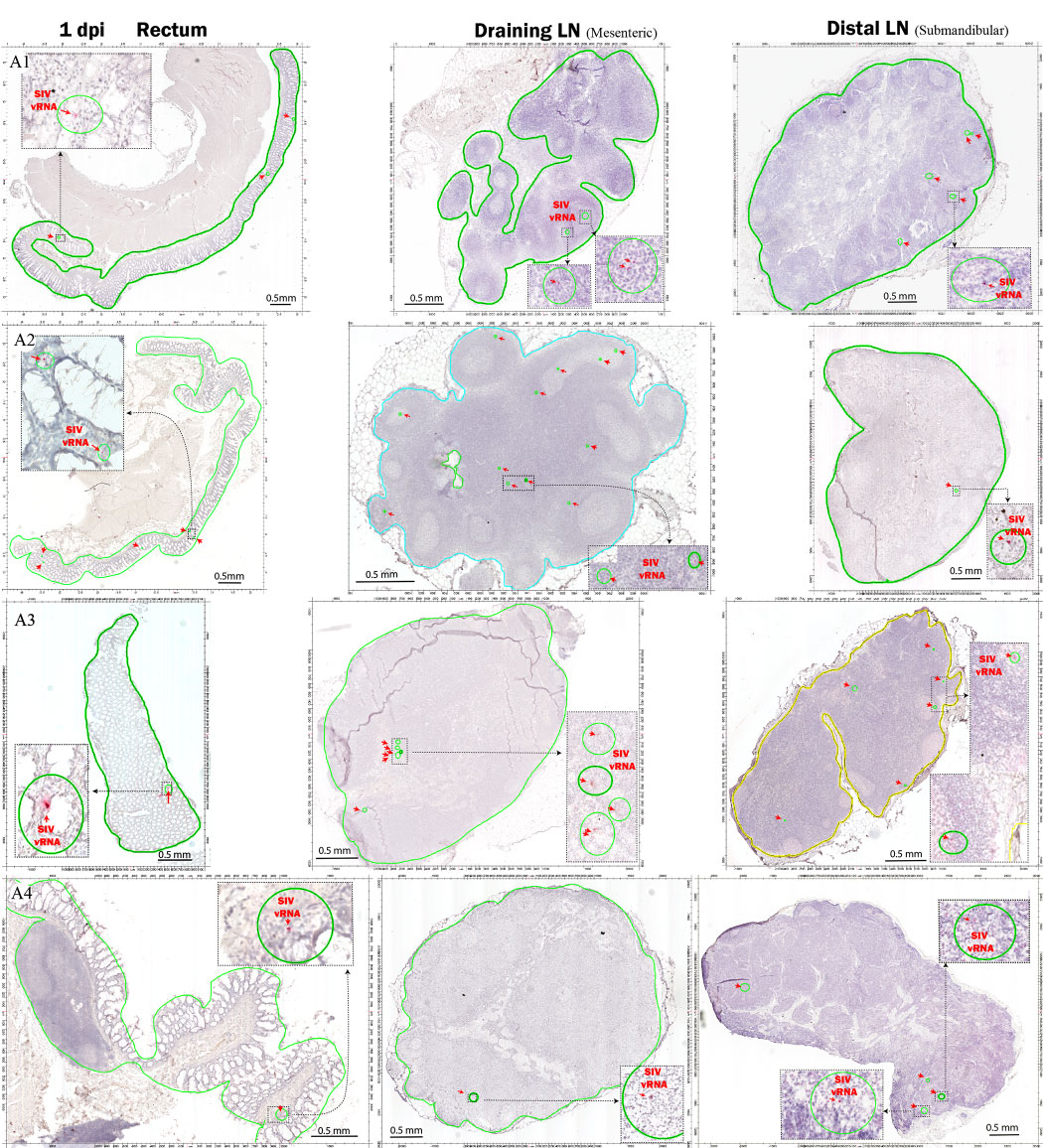

### Supplemental Figure 3

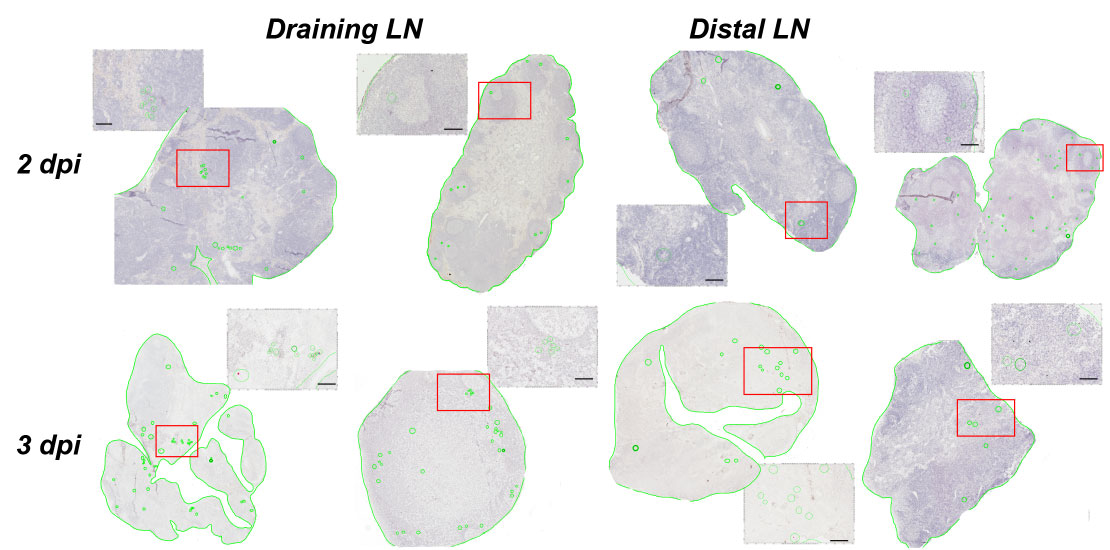

### Supplemental Figure 4

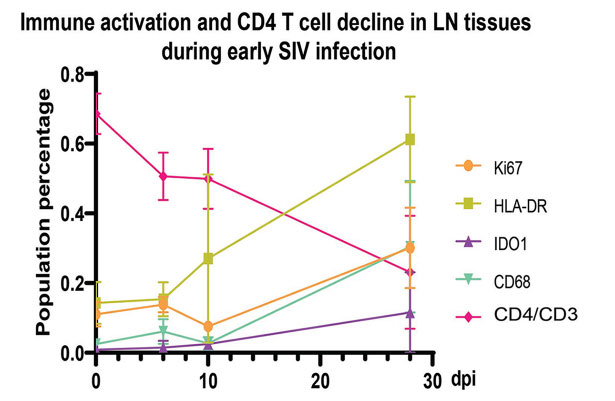

### Supplemental Figure 5

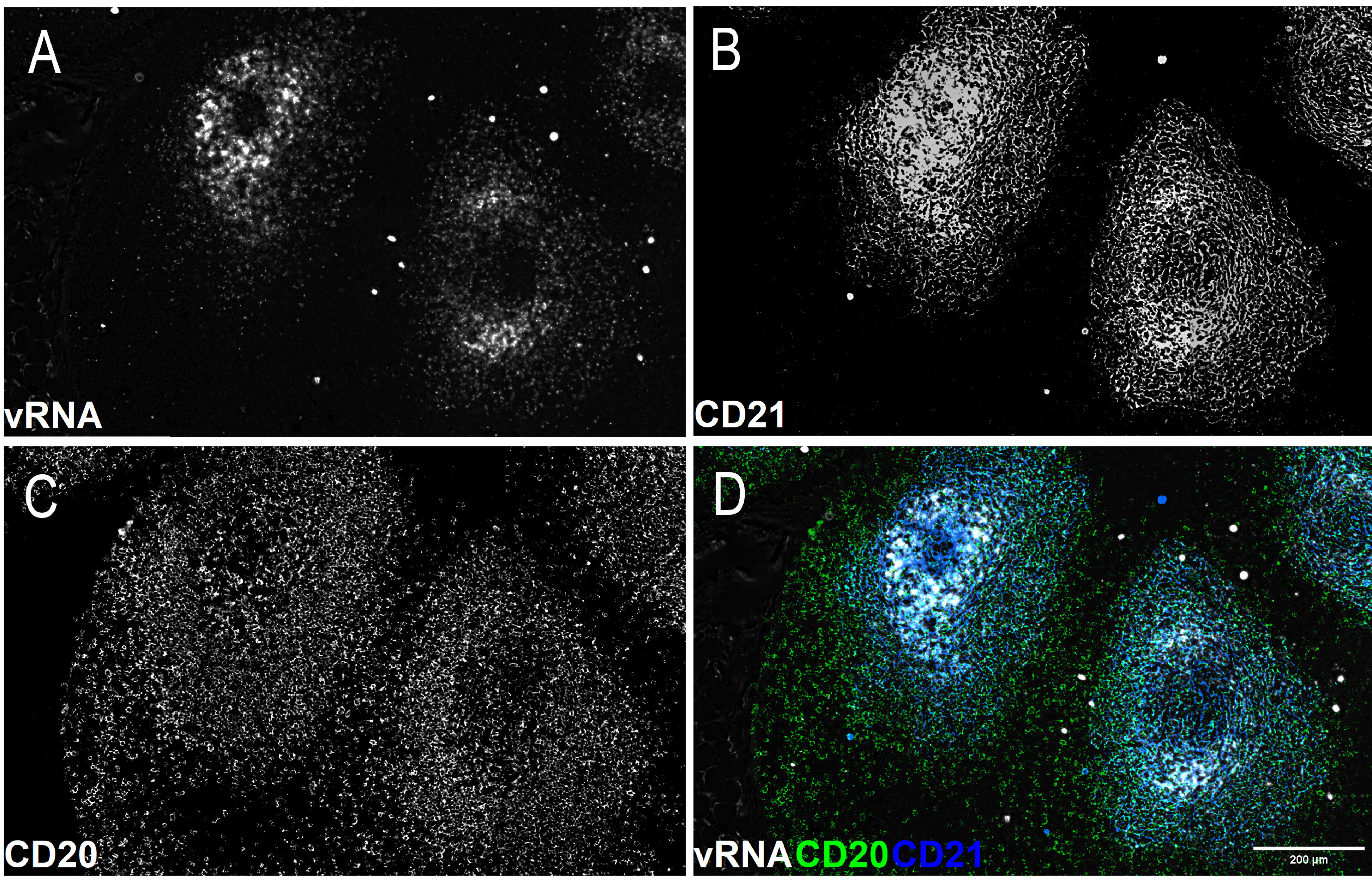

### Supplemental Table 1

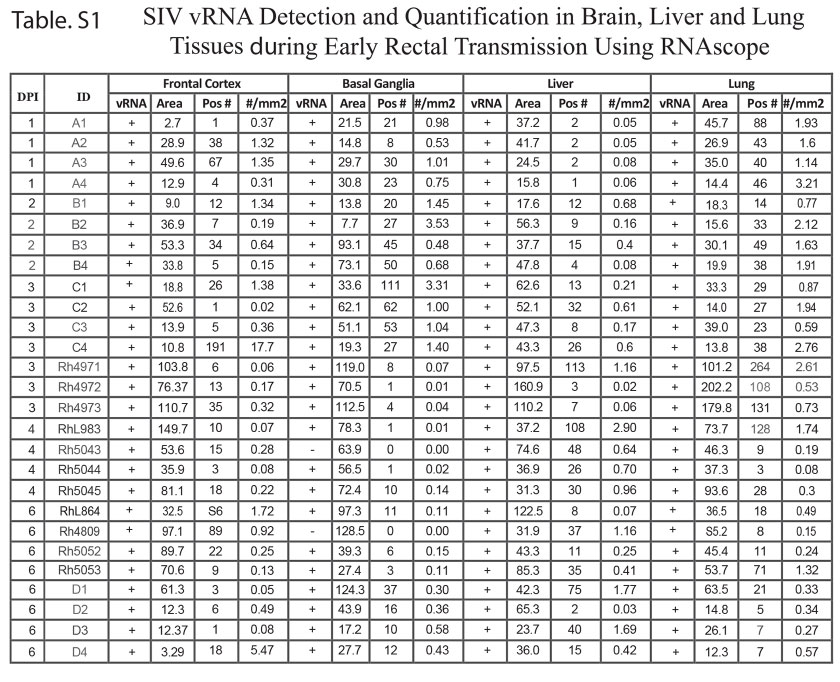
